# Divergent resistance mechanisms to immunotherapy explains response in different skin cancers

**DOI:** 10.1101/2020.09.15.298182

**Authors:** Emmanuel Dollinger, Daniel Bergman, Peijie Zhou, Scott Atwood, Qing Nie

## Abstract

The advent of immune checkpoint therapy for metastatic skin cancer has greatly improved patient survival. However, most skin cancer patients are refractory to checkpoint therapy, and furthermore the differential intra-immune signaling between cancers driving the response to checkpoint therapy remains uncharacterized. When comparing the immune transcriptome in the tumor microenvironment of melanoma and basal cell carcinoma (BCC), we found that the presence of memory B cells and macrophages negatively correlate when stratifying patients by response, with memory B cells more present in responders. Inhibitory immune signaling is reduced in melanoma responders and is increased in BCC responders. We further explored the relationships between macrophages, B cells and response to checkpoint therapy by developing a stochastic differential equation model which qualitatively agrees with the data analysis. Our model predicts BCC to be more refractory to checkpoint therapy than melanoma. We show differences in tumor progression and regression that could serve as a diagnostic and predict the optimal ratio of macrophages and memory B cells for successful treatment.

## Introduction

Checkpoint immunotherapy can drive durable responses in many metastatic cancers, with most adverse events being grade 1 or 2 ^1–4^. Current FDA-approved checkpoint inhibitors fall into two categories: CTLA-4 inhibitors and PD-1/PD-L1 inhibitors. CTLA-4 expressed by T regulatory cells (Tregs) outcompete costimulatory molecules on cytotoxic T lymphocytes (CTLs) necessary for their activation, which results in anergy and eventual apoptosis. Cancer and immune cells express PD-L1, which binds to PD-1 expressed by effector cells including CTLs and also by innate immune cells such as NK cells^5^. Binding of PD-1 on effector cells inhibits their cytotoxicity and also promotes anergy and eventual apoptosis ^1^. Inhibition of either pathway leads to durable cancer regression in many cancers with varied somatic mutations ^1,3^. Checkpoint therapy’s utility remains limited, however, with most patients either not responding or acquiring resistance to treatment ^3^.

Despite the promise of cancer checkpoint immunotherapy, our understanding of how these therapies affect a system as responsive and dynamic as the immune system remains incomplete. Many studies focus on the effect of checkpoint therapy on CTLs ^1,6–10^. Notably, two major recent studies sequenced the transcriptome of the tumor microenvironment (TME) at a single-cell level before and after checkpoint therapy in melanoma ^9^ and in basal cell carcinoma (BCC) ^8^, and both these studies focused on the effect of checkpoint therapy on CTLs. However, effects of checkpoint therapy on different immune cell types have been previously observed ^10–12^. In a phase I clinical trial for nivolumab, divergent and even opposite effects of nivolumab on T cells and B cells were observed ^11^. More recently, B cells have been shown to correlate with response to checkpoint immunotherapy even more strongly than CTL presence ^10,12^; however, this remains contentious, with other studies showing no effect ^13^.

Since the potential of single cell RNA-sequencing (scRNA-seq) was demonstrated for the first time on blastomeres in 2009^14^, the ability to partially capture the transcriptome of individual cells has driven insight in many disparate areas of research, including understanding myoblast differentiation ^15^, identifying rare cancer populations^16^, and others^17,18^. scRNA-seq is particularly well-suited to holistically analyze different immune cell types, due to its ability to capture high-resolution transcriptomic data from many cell types at once. Dynamical systems modeling has previously been used to model the TME^19–21^ (reviewed in ^22^) and can be parameterized by scRNA-seq data analyses to explore roles of regulations and predict responses to immunotherapy, pointing ways to new therapeutic interventions.

To compare and contrast the immune responses of responders and non-responders to checkpoint therapy, we analyzed scRNA-seq datasets from BCC and melanoma ^8,9^ and found that memory B cells are most present in responders and vis-versa for macrophages. We characterized their cellular signaling and found that macrophages strongly inhibit memory B cells in melanoma nonresponders. However, the immune inhibitory signaling increases in responders in BCC nonresponders, along with a strong increase in PD-1 signaling. To fully explore the dynamics of the system, we built a three-state dynamical continuum model that predicts the responsiveness to immunotherapy rests on a low number of B cells and a high number of macrophages pretreatment. In addition, the model predicts a patient’s tumor burden in conjunction with the number of memory B cells in the TME at any point after treatment with checkpoint therapy is sufficient to determine whether the patient will have a durable response.

## Results

### BCC and melanoma exhibit similar responses to checkpoint immunotherapy

To characterize differences between responders and non-responders after checkpoint immunotherapy, we analyzed two scRNA-seq datasets from melanoma ^9^ and BCC ^8^ patients before and after immunotherapy. The melanoma scRNA-seq dataset consists of 48 FACS-sorted CD45+ samples from 32 patients with metastatic melanoma before and after either anti-PD-1, anti-CTLA-4, or combination treatment. The BCC dataset consists of 24 site-matched samples from 11 patients with metastatic or locally advanced BCC before and after PD-1 blockade. We clustered the immune cells from each dataset separately and found they both contain CD4+ T cells, CD8+ T cells, T regulatory cells (Tregs), macrophages, memory B cells, plasma B Cells, plasmacytoid dendritic cells (pDCs), and cycling cells (**Figure 1 A-B; Supplementary Figure 1**). Overall our clustering recapitulated the original analysis ^8,9^ (**Supplementary Figure 1)**. We then calculated the percentage of immune cells by responders or non-responders and found that the overall percentage of responders and non-responders in each cluster was similar across both cancers (**Figure 1C**). Between BCC and melanoma, CD8+ T cells consistently showed roughly equal distribution between responders and non-responders, whereas memory B cells were highly concentrated in responders and macrophages were highly concentrated in non-responders. To determine whether these trends are generalized or patient-specific, we compared the percentage of macrophages, memory B cells, and CD8+ T cells stratified per patient and separated by responders and non-responders (**Figure 1D-F**). We found that macrophages represent a higher percentage of cells per patient in non-responders and that the opposite is true for memory B cells. Overall, the distribution of immune cells in responders and non-responders show remarkable similarity in both cancers, despite the differences in immunogenicity and sequencing technologies used for each cancer.

**Figure 1:**
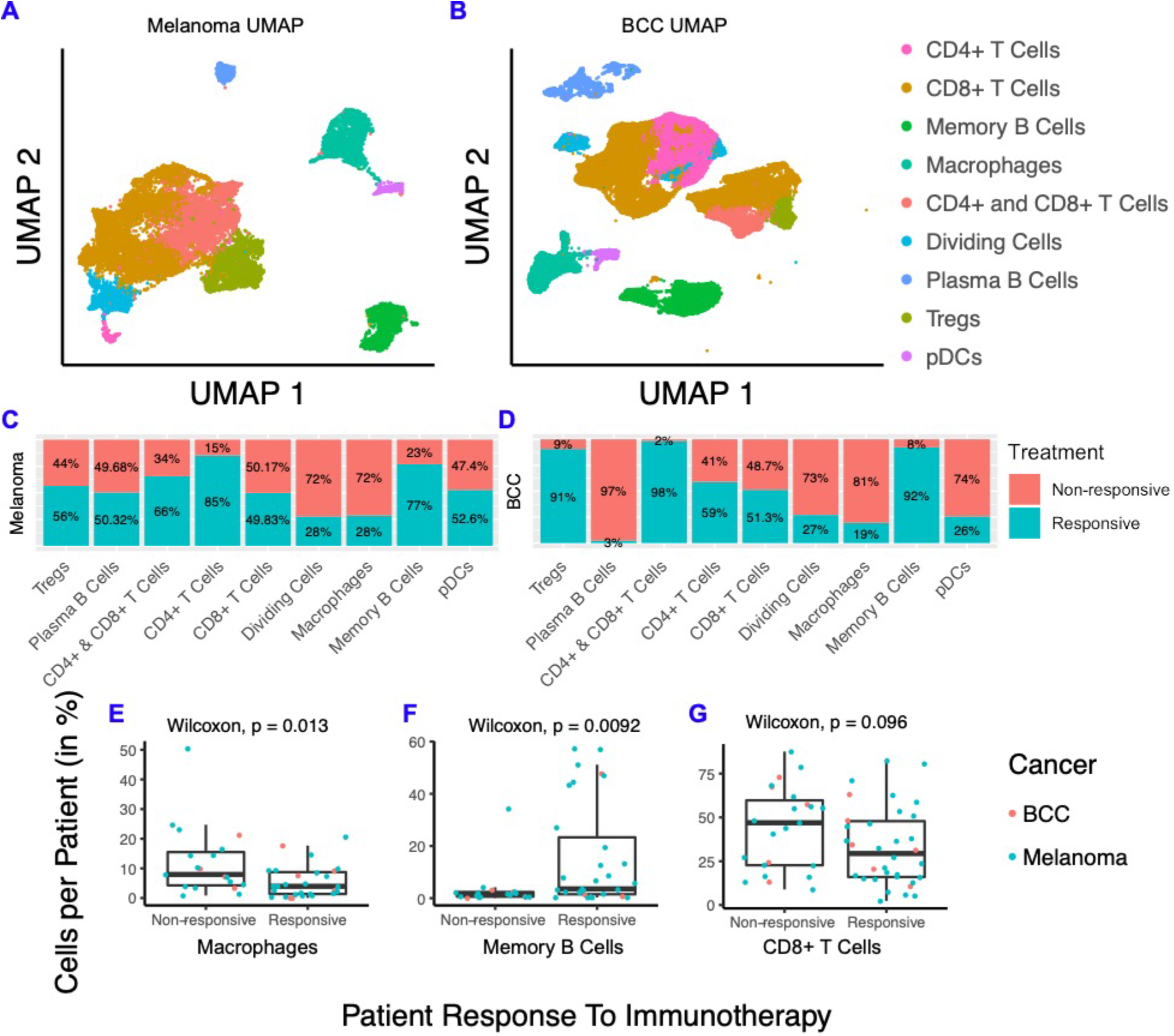
Melanoma and BCC have similar responses to immunotherapy. (A, B): Dimensionality reduction of melanoma (A) and BCC (B). (C) Distribution of cells from responders and non-responders, grouped by cluster. (D-F): Percentage of macrophages (D), memory B cells (E) and CD8+ T cells (F) per patient, grouped by responders and non-responders.

### Memory B cells are more active in post-treatment responders and anergic in posttreatment non-responders

As memory B cells were highly concentrated in immunotherapy responders in both datasets and may provide insight into mechanisms by which patients respond, we subclustered the memory B cells in both datasets and found the melanoma memory B cells to be well-mixed with regards to treatment, response, and patient, whereas the BCC memory B cells suffer from batch effects stemming from the small patient size (**Figure 2A-B; Supplementary Figure 2**). When comparing the memory B cell subclusters between BCC and melanoma, we observe differences in gene expression unique to each cancer (**Supplementary Figure 2C**) suggesting the memory B cells are not occupying similar states and may be differentially interacting with the TME.

**Figure 2:**
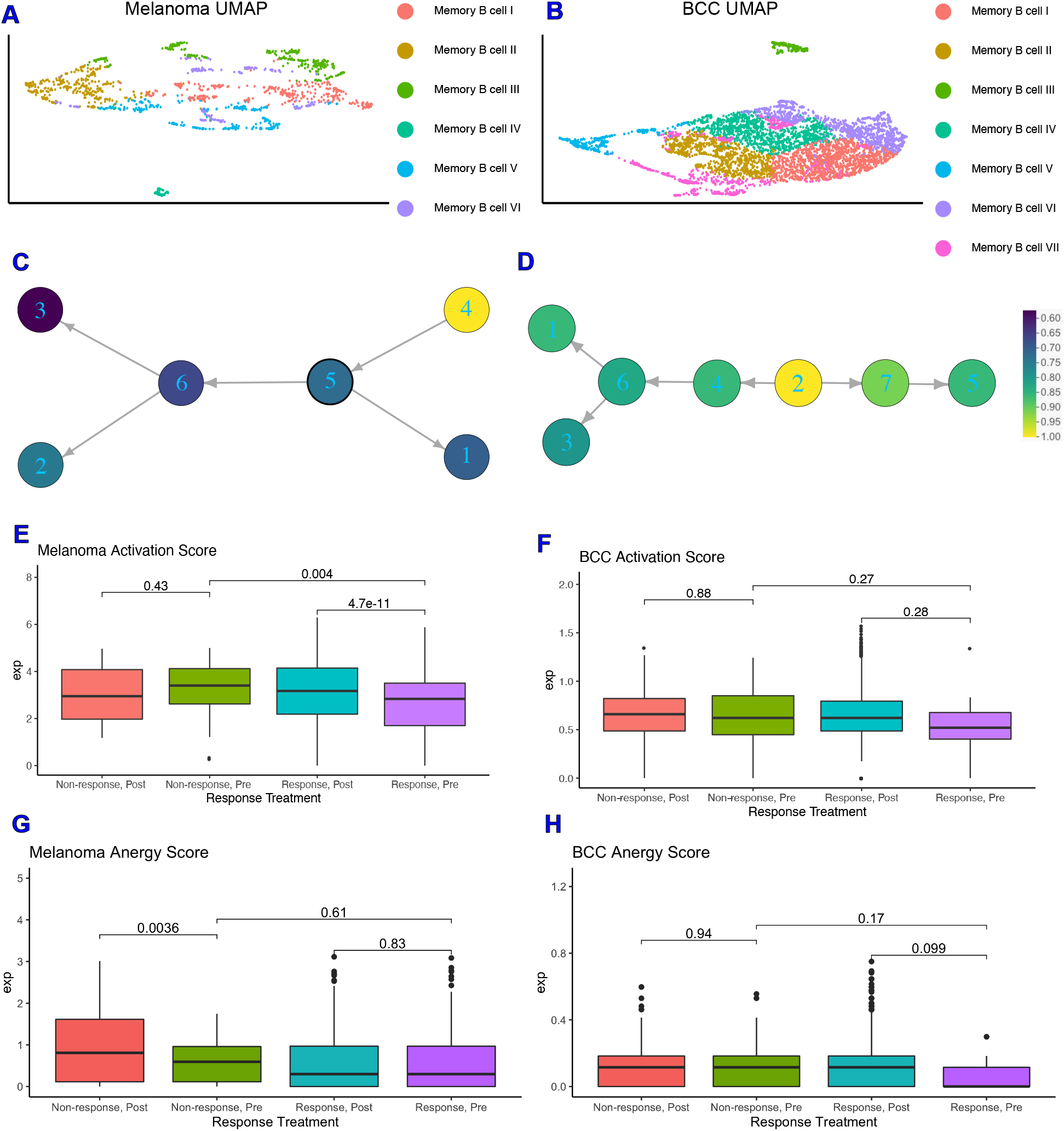
Memory B cells are more activated in non-responders pre response. (A, B) Dimensionality reduction of the memory B cells subsets of melanoma (A) and BCC (B). (C, D) Psuedotime ordering of melanoma (C) and BCC (D), colored by normalized activation score within each dataset. (E, F) Activation scores of memory B cells in melanoma (E) and BCC (F). (G, H) Anergy scores of memory B cells in melanoma (G) and BCC (H).

With the increase in Memory B cell complexity, we used similarity matrix-based optimization method (SoptSC) to infer their lineage ^23^.The melanoma and BCC memory B cell lineages show distinct trajectories that reflect the differences in cellular states between the two cancers (**Figure 2C-D**). However, when segregating the pseudotime trajectory along activation scores that reflect memory B cells binding to their specific antigen and actively expressing costimulatory receptors for T helper 1 (Th1) cells ^24^, both lineages show an increase in activation score at their terminus (**Figure 2C-D**). Both responders and non-responders show the increase in activation at the trajectory terminus, suggesting that the immune system is attempting to activate memory B cells in distinct ways within each cancer.

To further define how memory B cells are interacting with their environment, we developed a score for memory B cell anergy to go along with the activation score (**Supplemental Figure 2**). If activated B cells don’t receive costimulatory signals from Th1 cells, they become anergic, non-responsive to stimulation, and eventually apoptose ^24^. The average normalized expression of each set of genes that make up activation or anergy scores were calculated for each cell and stratified on pre- or post-treatment and response of the patient. In the melanoma dataset, the activation score for pre-treatment responders is significantly lower than in post-treatment responders, and the activation score is significantly higher in pre-treatment non-responders than in pre-treatment responders (**Figure 2E**), which makes sense given memory B cells should be more active in responders after treatment and not in non-responders. The only significant difference in the anergy scores for melanoma patients comes from the post-treatment nonresponders, which are more anergic than those in pre-treatment (**Figure 2G**). BCC memory B cells show similar, but not significant, trends in activation and anergy to memory B cells in melanoma (**Figure 2F, H**).

### Macrophages in BCC have a pro-inflammatory genotype, regardless of responder status

Macrophages have important roles in cancer immune suppression and correlate with poor prognosis, with drugs being developed to inhibit their suppressive ability ^25,26^. In particular, inhibiting the myeloid growth factor pathway CSF1/CSF1R increases the sensitivity of pancreatic ductal adenocarcinoma to checkpoint immunotherapy by decreasing the number of macrophages in the TME, increasing antigen presentation on macrophages, and increasing checkpoint ligands on tumors ^27^. To characterize the role of macrophages in resistance to immunotherapy in each cancer, we subclustered the macrophages and found that macrophages are mostly present in non-responders in both cancers (**Figure 3A-D; Supplementary Figure 3**). Similar to memory B cells, when comparing the macrophage subclusters between BCC and melanoma, we observe differences in gene expression unique to each cancer (**Supplementary Figure 3**) suggesting the macrophages are not occupying similar states and may be differentially interacting with the TME.

**Figure 3:**
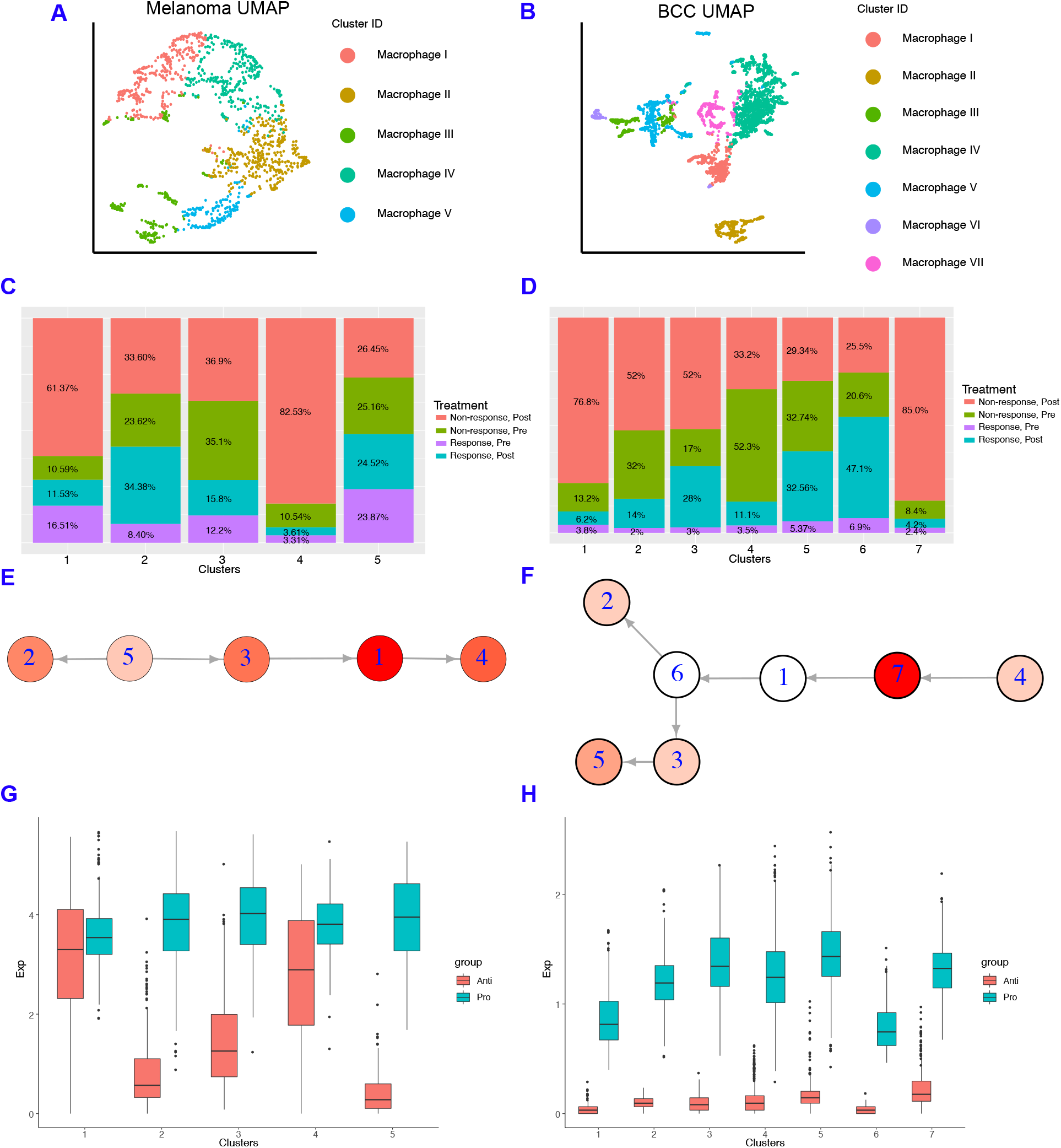
Macrophages in BCC have more of a pro-inflammatory genotype, regardless of responder status. (A, B) Dimensionality reduction of the macrophage subsets of melanoma (A) and BCC (B). (C, D) Percentage of responders/non-responders in pre/post treatment per macrophage cluster in melanoma (C) and BCC (D). (E, F) Average expression of anti- and pro-inflammatory genes by cluster of macrophages in melanoma (E) and BCC (F). (G, H) Psuedotime of macrophage clusters in melanoma (G) and BCC (H). Each psuedotime node is qualitatively colored by normalized expression of anti-inflammation score. The melanoma psuedotime correlates well with the percent of non-responders post-treatment, whereas the BCC psuedotime does not.

We built macrophage inflammatory scores using genes that are either “pro-inflammatory” or “antiinflammatory” (**Supplementary Figure 3**)^28,29^. In the melanoma dataset, we found that antiinflammatory gene expression correlates well with the percentage of macrophages found in post-treatment non-responders, indicating that macrophages in the melanoma TME are involved in the refractory response to immunotherapy. (**Figure 3E**). However, BCC macrophages have very low expression of anti-inflammatory genes in all backgrounds, suggesting that BCCs may not regulate immunotherapy response by inflammatory signals (**Figure 3F**). Although both cancers have more macrophages in non-responders, the macrophages have unique inflammatory signatures that are linked to different processes (**Supplementary Figure 3**). Using SoptSC to generate a lineage for macrophages, we observed distinct trajectories that reflect the differences in cellular states between the two cancers but a similar increase in anti-inflammatory scores at the trajectory terminus (**Figure 3G-H**).

### Anti-inflammatory signaling is reduced in melanoma responders and increased in BCC responders

To quantify changes of intra-immune signaling between B cells and macrophages with immunotherapy response, we used SoptSC to construct probabilistic cell-cell signaling interactions. Signaling probabilities are quantified based on the weighted expression of signaling pathway components between sender-receiver cell pairs inferred through expression of ligand-receptor pairs and their downstream targets (**Methods**)^23^. We subsetted memory B cells, plasma B cells, and macrophages in both datasets and calculated the probability of cluster-cluster cell signaling, which averages individual cell signaling probabilities within each cluster (**Figure 4A, E**). We included the plasma B cells in the analysis because of the stark difference in fraction of responders and non-responders between the two cancers (**Figure 1C**). We chose three pathways to study: FCGR2B, IL6, and PD-1. FCGR2B is a well-characterized inhibitory pathway used by macrophages to inhibit B cells^30^. The IL6 pathway has been correlated with B regulatory cell (Breg) activation, which has been implicated in many immunological tolerance mechanisms such as organ transplantation^31^, cancer^32^, and self-stimulation of tumor cells ^33^. The PD-1 pathway is used as a control for response.

**Figure 4:**
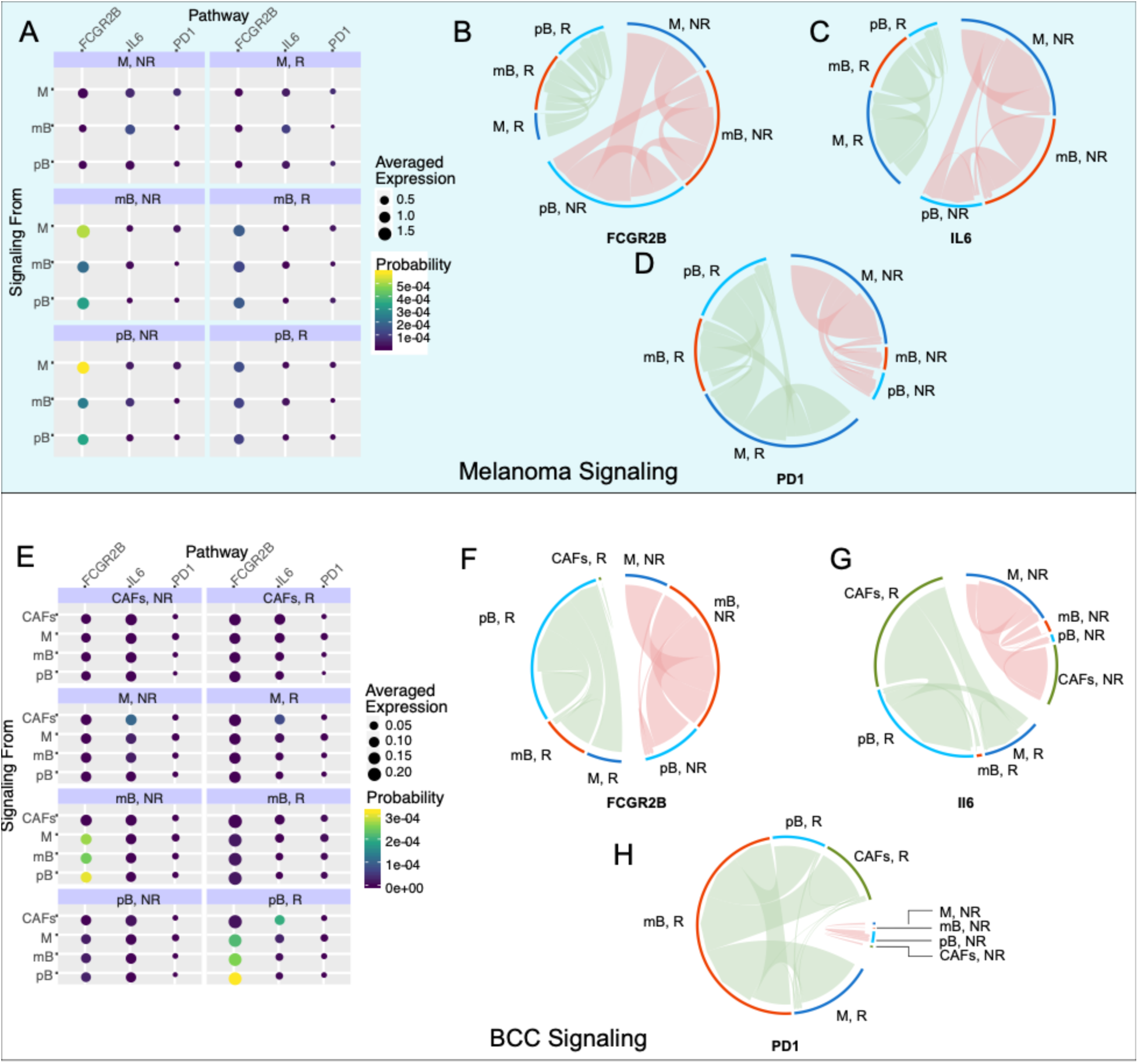
Inhibitory signaling is diminished in melanoma responders, whereas BCC responders experience increased inhibitory signaling. (A) and (E): Probability of signaling and averaged expression of the ligand/receptor/downstream target in each cell population for melanoma (A) and BCC (E). (B-D): Signaling of the FCGR2B (B), IL6 (C), PD1 (D) pathways for melanoma. (F-H): Signaling of the FCGR2B (F), IL6 (G), PD1 (H) pathways for BCC.

In the melanoma dataset, we found that the FCGR2B pathway is strongly upregulated in non-responders **(Figure 4B)**. The majority of the FCGR2B-mediated inhibition goes from macrophages to memory and plasma B cells, suggesting that B cells are selectively inhibited in non-responders. Concurrently, the IL6 pathway is upregulated in non-responder memory B cells, with signaling directed towards the macrophages and plasma B cells **(Figure 4C)**, suggesting an anti-inflammatory response in these cells and further suppression in immune response. PD-1 signaling is increased in responders, with the source of signaling switching from macrophages in non-responders to plasma B cells in responders **(Figure 4D)**.

In the BCC dataset **(Figure 4E)**, we subsetted macrophages, both B cell types, and cancer-associated fibroblasts (CAFs), which have high IL6 signaling. FCGR2B occurs with similar strength in responders and non-responders, with a switch from memory B cell inhibition in nonresponders to plasma B cell inhibition in responders **(Figure 4F)**. IL6 signaling increases in responders, with the majority of the signaling coming from CAFs and going to plasma B cells in responders **(Figure 4G)**. In non-responders, the signaling still comes from CAFs but now signals mainly to macrophages **(Figure 4G)**. The upregulation of IL6 with checkpoint blockade was previously characterized in melanoma mice models but not to this current resolution ^34^. Finally, the PD-1 signaling is drastically upregulated in responders, with the majority of the signaling directed towards memory B cells in BCC instead of plasma B cells in melanoma **(Figure 4H)**. These results, especially the trends of the FCGR2B and IL6 pathways, indicate that the immune system in melanoma is actively inducing an immune suppressive environment, which is contributing to resistance; however, BCC seems to only be inducing a suppressive environment in responders, indicating that there is a different mechanism of resistance, relying on the simple lack of sufficient activation of immune cells during therapy.

### A dynamical model on interactions among memory B cells, macrophages, and skin tumors

To better understand the dynamics of the immune system during treatment and specifically predict the best immune cell composition for response, we developed a three-state continuum dynamical model based on the bioinformatic clustering, lineage, and signaling analyses. We chose cancer, B cells, and inhibitory macrophages (referred to as simply “macrophages”) as our state variables (**Figure 5A** and **Methods**). The cancer undergoes logistic growth and has four possible steady states: none, low (~10^3^ cells), high (~10^8^ cells), and very high (~10^9^ cells). The B cells kill cancer cells and macrophages inhibit B cell proliferation. The parameters of the dynamical model were selected based on our bioinformatic analyses and previous literature (**Supplementary Table 1**), and the equations for the dynamical model are shown in **Methods**.

**Figure 5:**
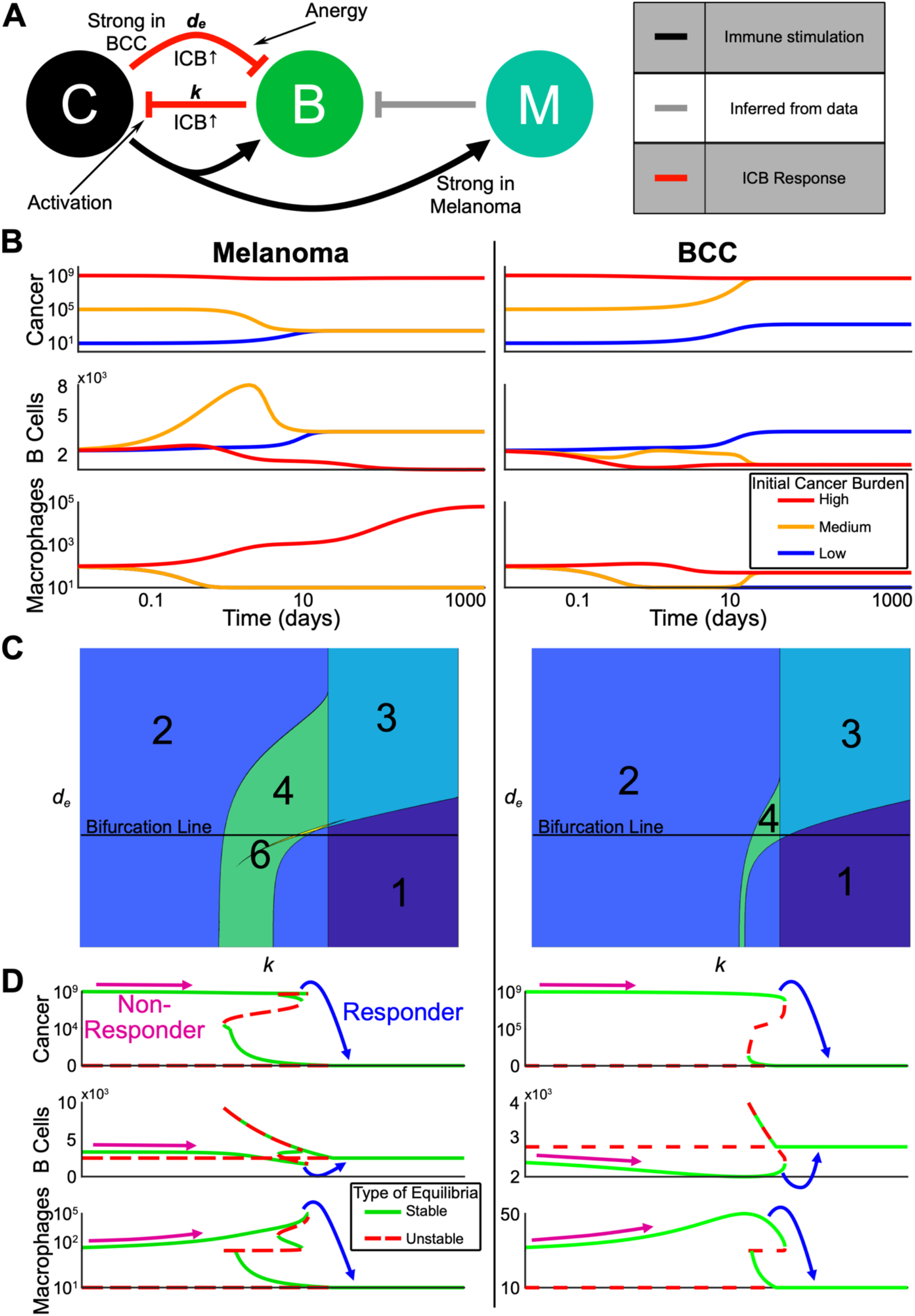
Multi-stability from complex interactions in melanoma and BCC can explain heterogeneous response to ICB. (A) Schematic representation of the model (C=cancer, B=B cells, M=anti-inflammatory macrophages). Red arrows indicate processes assumed to be upregulated by ICB. Grey arrow is inferred from our single-cell analysis. (B) Three trajectories with varying initial cancer population for each of melanoma and BCC. Color of the trajectory corresponds to initial cancer population. All axes are log scale. (C) Contour plots showing varying number of equilibria as the death rate of B cells (y-axis) and killing rate by B cells of the cancer (x-axis) are varied. The “Bifurcation Line” indicates the values for which the bifurcations in Panel D are plotted. (D) Bifurcation diagram in the killing rate, *k*, corresponding to the Bifurcation Line in Panel C. Possible starting and ending values for a non-responder and a responder are shown. Each Cancer and Macrophage axis is shown over two different scales for visualization purposes.

We made two assumptions from the signaling analyses that differentiate the divergent refractory mechanisms of each cancer. First, consistent with macrophages in BCC having less of an anti-inflammatory phenotype (**Figure 3F**), we assumed that the cancer-mediated up-regulation of macrophage proliferation is weaker in BCC relative to melanoma. Second, consistent with stronger B cell suppression in BCC (**Figure 4B-C, F-G**), we increased the negative regulation of B cells by BCC cancer cells relative to that in melanoma (all parameters are shown in **Supplementary Table 1**).

To understand the possible dynamics predicted by the model without immunotherapy, we computed several representative trajectories (**Figure 5B**). We fixed the starting immune populations and each parameter (**Supplementary Table 1**) and varied the initial cancer burden between low, medium, and high. In both melanoma and BCC, the high cancer burden remained high while the immune populations followed different trajectories. Macrophages steadily increased in melanoma and remained low in BCC in accordance with previous observations ^35,36^. Both melanoma and BCC were not able to transition from a low cancer burden to a high one in the chosen set of parameters. The medium cancer burden regressed to a low cancer burden only in melanoma while accompanied by a transient spike in memory B cells, whereas the medium cancer burden in BCC progressed to a high cancer burden.

### The model predicts the most likely immune cell composition for responders and shows BCC is less likely to response to treatment

To understand the effects of immunotherapy on cancer burden, we analyzed the steady states of our model within certain biologically relevant parameter ranges (**Supplementary Table 1**). Overall, our system displayed multi-stability, a common concept in cancer state modeling^19^, in both melanoma and BCC: the system could evolve towards two or more steady states depending on the level of cancer burden (**Figure 5C**).

We decided to vary the killing rate of B cells *k* as a proxy for immunotherapy, and study how the steady states change as we increased *k* (i.e. bifurcation analysis). We found that melanoma and BCC responded similarly to immunotherapy (**Figure 5D**). We observed responders in both cancer backgrounds where a very high cancer burden transitioned to a low cancer burden. In melanoma responders, an increase in B cells and a large decrease of macrophages was observed. We saw the same pattern in the immune profile of BCC, except the increase in B cells was larger while the decrease in macrophages was smaller. On the other hand, non-responders showed a small decrease in B cells and an increase in macrophage population, potentially up to several orders of magnitude in the melanoma case.

To predict responsiveness to immunotherapy, we determined the immune cell composition for responders and non-responders pre-treatment. We compared the equilibrium number of macrophages and B cells just before and just after the transition from non-responders to responders as we increased the B cell killing rate *k* (**Figure 5D**). Compared to non-responders, responders pre-treatment had a lower B cell population and a higher macrophage population, which strongly decreases post-treatment.

In our chosen parameter regime, the value of the immunotherapy killing rate at which a patient would become a responder was lower for melanoma than BCC (**Figure 5D**). This relationship between the two cancers persisted even as we varied *d_e_* (i.e. the death rate of B cells) leading us to predict melanoma to be more likely to respond to immunotherapy than BCC (**Supplementary Figure 4**).

### Noise-induced cancer progression and regression potentially account for therapy-resistance in BCC

In the highly complex cancer-immune interacting environment, fluctuations in cell populations may induce random transitions among meta-stable states ^21,37^. We therefore incorporated stochastic effects into our three-component dynamical model (equations detailed in **Methods**). In our stochastic model, the inclusion of random fluctuations in cell population dynamics allows for spontaneous (as opposed to by varying a parameter) transitions between cancer states with various burdens, contributing another source to affect the checkpoint therapy outcome by the spontaneous progression or regression of cancer ^38–40^.

In order to compare the relative stability of noisy cancer states, we constructed a cancer-state landscape to visualize the global structures of attractor basins in melanoma and BCC populations and their transition dynamics (**Figure 6A**). The less likely cancer states correspond to shallower basins in the landscape – the intuition here is analogous to the classic Waddington landscape for cell fate commitment^41^, or wells in activation energy barrier diagrams. The deeper the well, the higher the energy required and the less likely the transition to a different well becomes. The cancer-state energy landscape agrees with our bifurcation analysis by showing two connected energy wells representing “stable” cancer states with “low” and “high” tumor burdens. The connectivity between these cancer wells suggests that spontaneous transitions can occur in both cancer types, corresponding to tumor progression and regression. We also observed that the cancer well of the low-burden state in BCC is shallower than in melanoma and the high-cancer state in BCC is deeper than in melanoma, suggesting a higher probability to transition to the higher-burden state and a smaller probability of the reverse transition (matching our prediction of BCC response from Figure 5) (**Figure 6A**).

**Figure 6:**
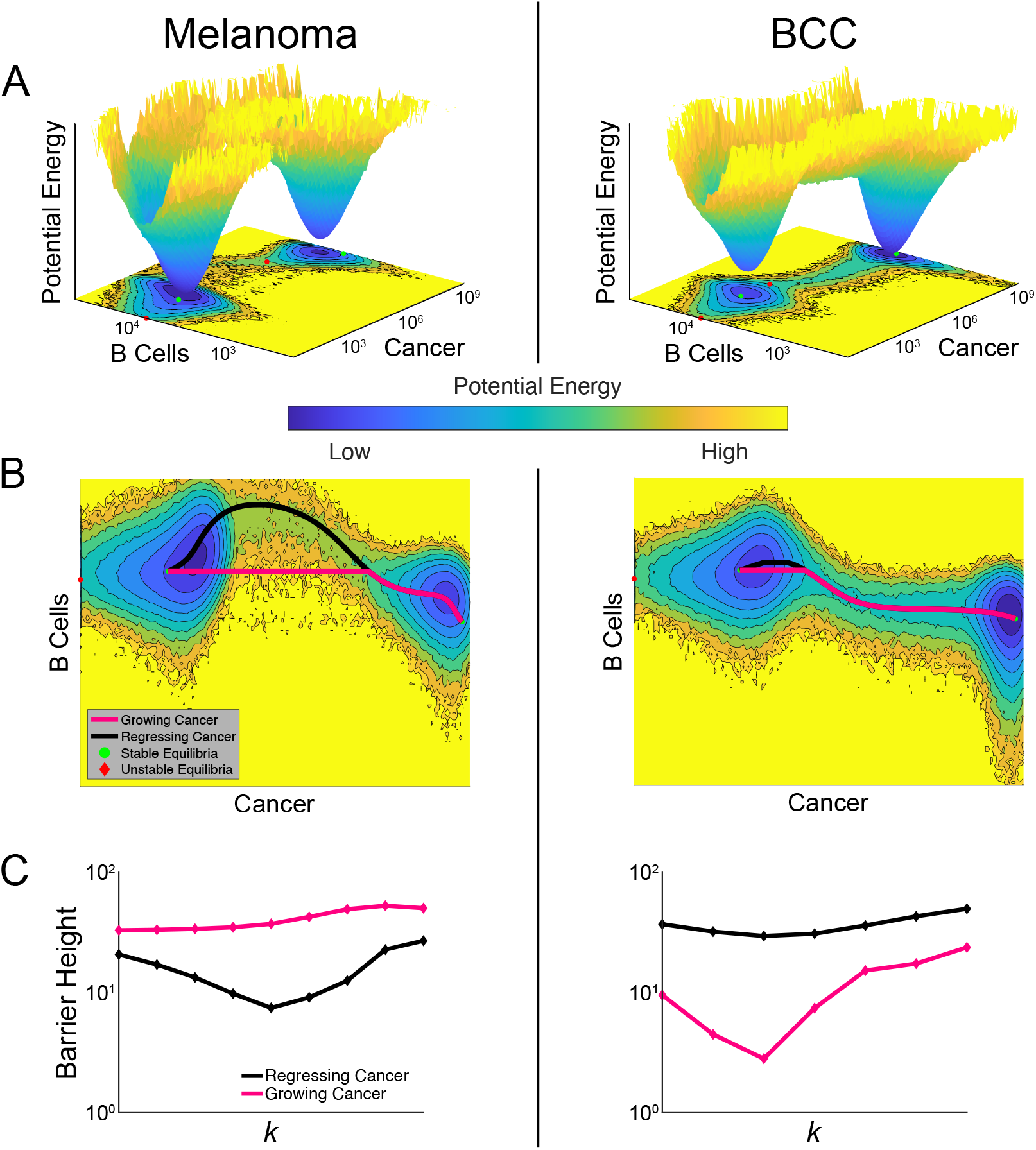
Comparisons of landscapes and transition paths for melanoma and BCC potentially explain the refractory response to ICB in BCC. (A) Cancer-state energy landscape of both cancers with *k* = 1.8 × 10^−4^ and *d_e_* = 1. Lower values indicate a higher probability of finding the system in that state. These values are the marginal probabilities having marginalized over macrophages. A contour plot is shown below along with the ODE-determined equilibria, both stable (green dots) and unstable (red diamonds). (B) The transition paths between the stable equilibria plotted over the contour plots from Panel A. The black path is for a regressing cancer and the magenta path is for a growing cancer. (C) The barrier height between the two high-burden and low-burden cancer states as it varies with the killing rate *k*. The black curve shows the variation of the barrier height for a regressing cancer, and the magenta for a growing cancer.

A unique feature of stochastic vs deterministic (e.g. the model of Figure 5) systems is the possibility of a transition between stable states. The specific transition path the system follows can discriminate between a growing and regressing cancer. To study these transition paths, we implemented the geometric minimal action method (gMAM) which determines the likelihood of each path (**Methods**)^42^. When melanoma transitioned from a high cancer state to a low cancer state (i.e. regresses) there was a strong increase in B cells, which was not true of the reverse transition (**Figure 6B**). In BCC, there is a similar pattern in the B cell population, though it is less pronounced.

To quantify how checkpoint therapy affects the likelihood of spontaneous tumor progression and regression, we calculated the change in activation energies between the two cancer states as the killing rate is increased (**Figure 6C**). Comparing these two curves, melanoma exhibited greater sensitivity to therapy with the activation energy decreasing more quickly. However, both cancers exhibit a surprising characteristic: the activation energy for regression initially decreases in the bistable region before growing at the higher end of this region. This indicated that a failure to push the system into a state with a unique attractor—a single, low cancer burden one—could make the cancer less likely to spontaneously regress. We found that the barrier height for regression in BCC is generally larger than in melanoma with similar killing rates, predicting BCC patients to be more refractory to immunotherapy in general.

When we quantified the activation energy for progression, we first observed that it was higher for melanoma than BCC, indicating a higher propensity for melanomas to have a durable response. We also noted that in BCC, the activation energy for progression is more sensitive to immunotherapy. At lower values of the killing rate *k*, therapy drove this barrier down making it more likely for an initial response to be reversed. This may provide a potential explanation for the unsatisfactory outcome of checkpoint therapy in BCC^39,40,43–45^. However, this trend eventually reverses and at higher killing rates, therapy makes spontaneous progression less likely.

## Discussion

Despite immunotherapy significantly advancing cancer therapy, not much is broadly known about the effects of immune cell interactions on patient response. We analyzed and compared two scRNA-seq datasets from melanoma and BCC and found that memory B cells are overrepresented in responders, whereas macrophages are over-represented in non-responders. We found that overall inhibitory signaling increased in melanoma non-responders and in BCC responders. The dynamical model that we constructed matched qualitatively with the data and predicted divergent responses to checkpoint therapy for responders and non-responders, as well as differences in immunotherapy response by BCC and melanoma.

Melanoma is a relatively rare and very dangerous immunogenic disease that arises from neural crest cells, whereas BCC is a very common and relatively benign non-immunogenic disease that arises from stem cells of the skin and hair follicle. However, our data suggests that their immune cell composition between responders and non-responders to immunotherapy is very similar, albeit for different reasons. Melanoma-associated macrophages in non-responders seem to be more anti-inflammatory, suggesting that macrophages may be an important resistance mechanism to immunotherapy as suggested in pancreatic cancer^27^. BCC-associated macrophages seem to be more pro-inflammatory, suggesting they are not important to immunotherapy resistance and that the barrier to BCC response to immunotherapy is a matter of immune cell recruitment and activation, not overcoming resistance. This matches well with reports that there is a sharp increase of immune cells after checkpoint therapy ^8^.

Our bifurcation analysis indicated that, dependent on the individual sensitivity toward the therapy in increasing the cancer killing rate *k* of B cells, the patient may either have a durable response, a partial response or a refractory response. These results could explain why some patients appear to have a naturally acquired resistance to immunotherapy ^46^. Our model suggests a low memory B cell count and high macrophage count (relative to each cancer) would indicate a likely response. This matches with the intuition that if there are less memory B cells, there was less of an immune response pre-treatment and the memory B cells would be less “exhausted”, matching with the results of Figure 2.

Despite the similarities in cell composition in each cancer, we also found important differences in the dynamics of melanoma and BCC cancers. From our energy landscapes, we observed a shallow low cancer burden well in BCCs, suggesting BCCs have a higher probability to transition to a higher cancer burden than melanoma. Our analysis of activation energies additionally suggests that BCC is less likely to respond to checkpoint therapy and the likelihood of posttherapy cancer recurrence is higher than in melanoma. BCC’s resistance to immunotherapy seems to be borne out in the literature, although more studies need to be done ^39,40,43–45^. In fact, the model suggests that an insufficient dose of immunotherapy could have adverse effects for some BCC patients with low pre-therapy killing rate, increasing their risk of tumor progression.

A crucial assumption we have made throughout this study is that memory B cells are directly affecting the cancer, either by releasing pro-inflammatory cytokines or by antibody production. Unfortunately, we were unable to verify whether these memory B cells were producing more antibodies in responders from the scRNA-seq datasets. Furthermore, it is unclear why memory B cells are more implicated in this response than plasma B cells. Memory B cells are known to produce antibodies with higher affinities compared to plasma B cells^47^, but require periods where they are not stimulated to properly mature, perhaps implying that the level of activation of B cells in responders before treatment needs to be relatively lower, at least for a period of time. Indeed, this intuition matches well with our results that memory B cells in responders pre-treatment are less activated than in non-responders pre-treatment.

## Methods

### Clustering

All analyses unless otherwise noted was the same for both datasets. The UMI count matrix for ^9^ were provided by personal communication from the authors, and the UMI matrix from ^8^ were downloaded via GEO, accession GSE123813. We excluded all cells with counts less than 200. The UMI counts were normalized and scaled using SCTransform in Seurat v3^48,49^; briefly, gene expression was normalized by taking the residuals of a generalized linear model that fits the counts of each gene across cells to a “regularized” negative binomial regression, with covariate cell sequencing depth. In this GLM, the Pearson residuals are the scaled gene expression values and were used for downstream analyses. These scaled gene values were used as input to PCA. The resulting first 30 dimensions of the PCA were used to generate the UMAP projections, with default parameters. The 30 first dimensions of the PCA were also used to calculate the shared-nearest neighbor network, which was used to cluster the cells (the smart local moving algorithm and resolution = 0.3 was used for clustering for both datasets; all other parameters were left as default).

To identify clusters, differential expression on each cluster was performed (Wilcoxon Rank Sum test; aside from thresholding the minimum fraction of cells that need to express a gene for that gene to be included to 0.25, all parameters were set to Seurat default) and the resulting top 50 differentially expressed genes were supplied to EnrichR, a gene list enrichment analysis tool ^50^. Clusters were identified by holistically considering different datasets (e.g. Human Gene Atlas, Mouse Gene Atlas, ARCHS4 Tissues and ARCHS4 Cell-lines).

To facilitate comparison across datasets, the T cell clusters were grouped by expression of CD8+ and/or CD4+; Tregs were identified by Enrichr and FOXP3+ expression. The B cells in the melanoma dataset were identified by Enrichr (plasma B cells) and specific markers (MS4A1 and CD40 for memory B cells). The B cells in the BCC dataset were identified by expression of the top differentially expressed genes between the memory B cells and plasma B cells in the melanoma dataset (plasma B cells: MZB1, IGHGP, IGHG3, IGHG1; memory B cells: CD79A, CD19, BANK1, IGHM, MS4A1).

### Lineage analysis and cell-cell signaling inference

The lineage analysis and cell-cell signaling was performed in SoptSC ^23^. SoptSC is a similarity matrix-based method for inferring cell lineage and cell signaling. Briefly, SoptSC calculates a cell-cell similarity matrix *S* based off of a low-rank representation of the log-transformed UMI count matrix. Our specific procedure for inferring clusters and building the cell lineage graph did not deviate from that laid out in ^23^: the similarity matrix was computed and the clusters and the number of clusters were inferred. For the non-responder subset of macrophages, memory B cells and plasma cells from BCC patients (Figure 4 C, D and E), the memory B cell cluster was manually defined based on their identities in the full dataset.

SoptSC calculates the probability that cells are signaling given a user-defined pathway of {Ligand, Receptor, Downstream upregulated target}. The cluster-cluster signaling graphs were generated by calculating the weighted graph of the cell-cell graph from the probability of signaling. Three pathways were considered: {FCGR2B, CD79A, FAS} and {FCGR2B, CD79B, FAS} were considered (i.e. calculated separately, then averaged) for macrophage-specific inhibitory signals, {PDL1, PD1, BATF} and {PDL2, PD1, BATF} were considered for PD1 signaling, and {IL6, IL6R, FCGR2B} was considered for immune inhibition. The probabilities were calculated in SoptSC by only considering probabilities > 0.025 and the resulting probability matrix was visualized in the *circlize* v0.4.9 package. The probabilities are relative to the transcriptomic information in each dataset and can only be compared with probabilities in the same dataset.

### Heatmaps, Dotplot, Barcharts and Box-and-whisker plots

For each dataset, the macrophages were subsetted, imported into SoptSC and clustered as described above. The cluster labels of the subsetted Seurat object were redefined with the SoptSC clusters, and the heatmaps were generated by inputting the specified gene list in the DoHeatmap function of Seurat. The dotplot was generated by redefining the cluster labels of the subset Seurat object with the SoptSC cluster labels and using Seurat’s Dotplot function, with the specified genes as input.

Taking the cluster labels of either the original Seurat clusters (figure 1) or the cluster labels of SoptSC (figure 2), the percent of either responder status (figure 1) or percent of response/treatment (figure 2) per cluster was calculated by dividing the number of cells in each category by the total number of cells in the cluster. The percent of cells per patient was calculated by dividing the number of specified cells (e.g. macrophages) by the total number of cells for that patient (we excluded the patients that had none of the specified cells). The Wilcoxon Rank Sum test was performed using the stat_compare_means function in the *ggpubr* package v0.2.5, with defaults. The “pro”-inflammation and “anti”-inflammation scores were calculated by averaging the normalized gene expressions for each gene list per cluster. Note that these cluster labels were generated using SoptSC and imported into the Seurat object, see above.

The gene intersection heatmaps for **Supplementary Figures 2C** and **3C** were calculated by doing differential expression on each subset as before and calculating the fraction of genes that were present in each cluster.

### The three-component dynamical model

Based on the single-cell data analysis, we modeled the dynamics of B cells, macrophages and cancer cells populations, which are emergent from their complex interactions. The assumptions on interactions between B cell and cancer cells are derived from existing literatures (**Supplementary Table 1**). The inclusion of anti-inflammatory macrophages (“macrophages”) and their interactions with other cells constitutes the novel aspect of our work, as most previous work uses pro-inflammatory cells as a third state variable (e.g. ^20^). Derived from the single-cell data analysis, the macrophages act to down-regulate B cell proliferation, directly in opposition to the cancer-mediated upregulation of that very process. The macrophages are in turn influenced by the cancer and memory B cells by responding to the apoptotic signals from cancer cells as the memory B cells kill them ^51^.

The model can be expressed in ordinary differential equations (ODEs). We let *C, B*, and *M* stand for the state variables of cancer cells, memory B cells, and macrophages, respectively. These three variables are time dependent. Cancer cells have a proliferation rate *a* and carrying capacity *b*^−1^. B cells kill cancer cells at rate *k*. B cells have a constant influx at rate *s* and die at rate *d_e_*. In the presence of cancer, B cells are stimulated and proliferate at a maximal rate *b_e_*. The cancer mediates this via a Hill function with EC_50_ term *κ_e_*. Macrophages inhibit this proliferation with another Hill function with EC_50_ term *κ_m_*. On the other hand, the cancer can adversely affect the B cell population by encouraging their removal from the system. This happens at a maximal rate of *d_e_* and with EC_50_ term *κ_d_*. Finally, macrophages also have a source, *g*, and death rate, *d_m_*. Their proliferation can be stimulated by apoptosis of cancer cells as induced by B cell killing, occurring at a maximal rate, *p*, and with EC_50_ term *κ_a_*. See **Supplementary Table 1** for parameter values and sources.

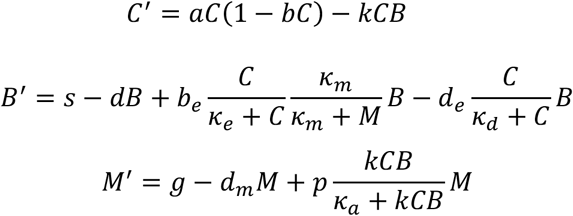

We conducted non-dimensionalization to simplify our analysis (**Supplementary Material**). To perform equilibria and stability analysis, we solved the derived 5-degree polynomial of steadystate equation and determine the stability using the eigenvalues of the Jacobian (**Supplementary Material**). The bifurcation plot can be generated by tracking the change of equilibria with respect to the parameter of interest.

To consider transitions among meta-stable cancer states, we included a time-independent noise term σ(X*_t_*) and generated a stochastic differential equation (SDE) model *dX_t_* = *b*(*X_t_*)*dt* + *σ*(*X_t_*)*dW_t_*, with *b*(*X_t_*) corresponding to the drift terms in the ODE system, and *W_t_* being a standard Weiner process.

### Cancer-state landscape and transition paths

The cancer-state landscape can quantify the relative stability of different meta-stable states perturbed by noise, closely relevant to the notion of energy landscape, a mathematical realization of Waddington’s epigenetics metaphor ^41,52,53^. To generate the landscapes, we simulated a large number of trajectories with randomly chosen initial conditions. Initial conditions were uniformly distributed over the log scale of the state variables. Each subsequent time step was binned based on the 3D coordinates and used to compute the probability a trajectory was in a particular bin. To arrive at the landscapes, we took the marginal probabilities over a given state variable and then computed the negative logarithm to arrive at our potential landscape.

To compute transition paths among meta-stable states, we applied the Freidlin and Wentzell’s (FW) large deviation theory ^42^, which states that under small noise assumption, the most probable path φ*(s) transiting from state *x*_1_ to *x*_2_ corresponds to the minimizer of the action functional

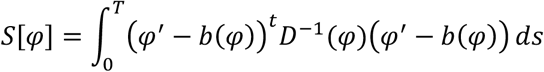

where matrix D(x) = σ(x)σ^t^(x) and 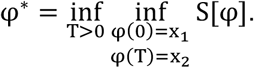.

We set *x*_1_ and *x*_2_ as stable fixed points of the ODE system. To tackle the numerical challenges introduced by critical points ^54^, we implemented a simplified geometric minimal action method (sgMAM) to solve the optimization problem ^54^. We used the action functional for these paths to compute the activation energies between stable equilibria. According to FW theory ^42^, the larger activation energies indicate longer mean transition time between metastable states.

## Supporting information

SFig 1

SFig 2

SFig 3

SFig 4

## Code and Data Availability

The data for ^9^ are stored in dbGAP phs001680.v1.p1, and the UMI matrix from ^8^ are stored in GEO, accession number GSE123813.

Code used to generate bioinformatic and mathematical results will be made available on Github and are available upon request.

## Author contributions

E.D., Q.N and S.A. conceived the project. E.D. and D.B. conducted the research and P.Z. contributed to the methods. Q.N. and S.A. supervised the research. E.D., D.B., P.Z., Q.N. and S.A. contributed to the writing of the manuscript.

## Acknowledgments

This work is partially supported by an NSF grant DMS1763272, a grant from the Simons Foundation (594598, Q.N.), and NIH grants P30AR075047, T32GM136624, U54CA217378 and R01CA237563, and a grant from the Hellman Fellowship. E.D. would like to thank the authors of ^9^ for the open and prompt sharing of their data. All authors declare no potential conflicts of interest.

## References

1. Ribas, A. & Wolchok, J. D. Cancer immunotherapy using checkpoint blockade. Science (80-.). 359, 1350–1355 (2018).

2. Topalian, S. L., Drake, C. G. & Pardoll, D. M. Immune checkpoint blockade: A common denominator approach to cancer therapy. Cancer Cell 27, 450–461 (2015).

3. Sharma, P., Hu-Lieskovan, S., Wargo, J. A. & Ribas, A. Primary, Adaptive, and Acquired Resistance to Cancer Immunotherapy. Cell 168, 707–723 (2017).

4. Linardou, H. & Gogas, H. Toxicity management of immunotherapy for patients with metastatic melanoma. Ann. Transl. Med. 4, (2016).

5. Liu, Y. et al. Increased expression of programmed cell death protein 1 on NK cells inhibits NK-cell-mediated anti-tumor function and indicates poor prognosis in digestive cancers. Oncogene (2017) doi:10.1038/onc.2017.209.

6. Postow, M. A., Callahan, M. K. & Wolchok, J. D. Immune checkpoint blockade in cancer therapy. J. Clin. Oncol. 33, 1974–1982 (2015).

7. Sharpe, A. H. Introduction to checkpoint inhibitors and cancer immunotherapy. Immunol. Rev. 276, 5–8 (2017).

8. Yost, K. E. et al. Clonal replacement of tumor-specific T cells following PD-1 blockade. Nat. Med. 25, 1251–1259 (2019).

9. Sade-Feldman, M. et al. Defining T Cell States Associated with Response to Checkpoint Immunotherapy in Melanoma. Cell 175, 998–1013.e20 (2018).

10. Helmink, B. A. et al. B cells and tertiary lymphoid structures promote immunotherapy response. Nature 577, 549–555 (2020).

11. Brahmer, J. R. et al. Phase I study of single-agent anti-programmed death-1 (MDX-1106) in refractory solid tumors: Safety, clinical activity, pharmacodynamics, and immunologic correlates. J. Clin. Oncol. 28, 3167–3175 (2010).

12. Petitprez, F. et al. B cells are associated with survival and immunotherapy response in sarcoma. Nature 577, 556–560 (2020).

13. Damsky, W. et al. B cell depletion or absence does not impede anti-tumor activity of PD-1 inhibitors. J. Immunother. Cancer 7, 1–7 (2019).

14. Tang, F. et al. mRNA-Seq whole-transcriptome analysis of a single cell. Nat. Methods (2009) doi:10.1038/nmeth.1315.

15. Trapnell, C. et al. The dynamics and regulators of cell fate decisions are revealed by pseudotemporal ordering of single cells. Nat. Biotechnol. (2014) doi:10.1038/nbt.2859.

16. Shaffer, S. M. et al. Rare cell variability and drug-induced reprogramming as a mode of cancer drug resistance. Nature (2017) doi:10.1038/nature22794.

17. Hwang, B., Lee, J. H. & Bang, D. Single-cell RNA sequencing technologies and bioinformatics pipelines. Experimental and Molecular Medicine (2018) doi:10.1038/s12276-018-0071-8.

18. Potter, S. S. Single-cell RNA sequencing for the study of development, physiology and disease. Nat. Rev. Nephrol. 14, 479–492 (2018).

19. Huang, S., Ernberg, I. & Kauffman, S. Cancer attractors: A systems view of tumors from a gene network dynamics and developmental perspective. Seminars in Cell and Developmental Biology (2009) doi:10.1016/j.semcdb.2009.07.003.

20. de Pillis, L. G. & Radunskaya, A. E. Modeling tumor–immune dynamics. in Springer Proceedings in Mathematics and Statistics (2014). doi:10.1007/978-1-4939-1793-8_4.

21. Huang, S. & Kauffman, S. How to escape the cancer attractor: Rationale and limitations of multi-target drugs. Seminars in Cancer Biology (2013) doi:10.1016/j.semcancer.2013.06.003.

22. Mahlbacher, G. E., Reihmer, K. C. & Frieboes, H. B. Mathematical modeling of tumor-immune cell interactions. Journal of Theoretical Biology (2019) doi:10.1016/j.jtbi.2019.03.002.

23. Wang, S., Karikomi, M., Maclean, A. L. & Nie, Q. Cell lineage and communication network inference via optimization for single-cell transcriptomics. Nucleic Acids Res. (2019) doi:10.1093/nar/gkz204.

24. B Cell Development, Activation and Effector Functions. in Primer to the Immune Response (2014). doi:10.1016/b978-0-12-385245-8.00005-4.

25. Palucka, A. K. & Coussens, L. M. The Basis of Oncoimmunology. Cell 164, 1233–1247 (2016).

26. Ruffell, B. & Coussens, L. M. Macrophages and therapeutic resistance in cancer. Cancer Cell 27, 462–472 (2015).

27. Zhu, Y. et al. CSF1/CSF1R blockade reprograms tumor-infiltrating macrophages and improves response to T-cell checkpoint immunotherapy in pancreatic cancer models. Cancer Res. 74, 5057–5069 (2014).

28. Ashburner, M. et al. Gene ontology: Tool for the unification of biology. Nature Genetics (2000) doi:10.1038/75556.

29. Carbon, S. et al. The Gene Ontology Resource: 20 years and still GOing strong. Nucleic Acids Res. (2019) doi:10.1093/nar/gky1055.

30. Smith, K. G. C. & Clatworthy, M. R. FcγRIIB in autoimmunity and infection: Evolutionary and therapeutic implications. Nature Reviews Immunology (2010) doi:10.1038/nri2762.

31. Peng, B., Ming, Y. & Yang, C. Regulatory B cells: The cutting edge of immune tolerance in kidney transplantation review-Article. Cell Death Dis. 9, (2018).

32. Sarvaria, A., Madrigal, J. A. & Saudemont, A. B cell regulation in cancer and anti-tumor immunity. Cell. Mol. Immunol. 14, 662–674 (2017).

33. Johnson, D. E., O’Keefe, R. A. & Grandis, J. R. Targeting the IL-6/JAK/STAT3 signalling axis in cancer. Nature Reviews Clinical Oncology (2018) doi:10.1038/nrclinonc.2018.8.

34. Tsukamoto, H. et al. Combined blockade of IL6 and PD-1/PD-L1 signaling abrogates mutual regulation of their immunosuppressive effects in the tumor microenvironment. Cancer Res. 78, 5011–5022 (2018).

35. Jiang, X. et al. Human keratinocyte carcinomas have distinct differences in their tumor-associated macrophages. Heliyon (2019) doi:10.1016/j.heliyon.2019.e02273.

36. Salmi, S. et al. The number and localization of CD68+ and CD163+ macrophages in different stages of cutaneous melanoma. Melanoma Res. (2019) doi:10.1097/CMR.0000000000000522.

37. Blank, C. U., Haanen, J. B., Ribas, A. & Schumacher, T. N. The ‘cancer immunogram’. Science (2016) doi:10.1126/science.aaf2834.

38. Kucerova, P. & Cervinkova, M. Spontaneous regression of tumour and the role of microbial infection – possibilities for cancer treatment. Anti-Cancer Drugs (2016) doi:10.1097/CAD.0000000000000337.

39. Sabbatino, F. et al. Resistance to anti-PD-1-based immunotherapy in basal cell carcinoma: A case report and review of the literature. J. Immunother. Cancer (2018) doi:10.1186/s40425-018-0439-2.

40. Lipson, E. J. et al. Basal cell carcinoma: PD-L1/PD-1 checkpoint expression and tumor regression after PD-1 blockade. J. Immunother. Cancer 5, 1–5 (2017).

41. Li, C. & Wang, J. Quantifying Cell Fate Decisions for Differentiation and Reprogramming of a Human Stem Cell Network: Landscape and Biological Paths. PLoS Comput. Biol. (2013) doi:10.1371/journal.pcbi.1003165.

42. Freidlin, M. I. & Wentzell, A. D. Random Perturbations. in (2012). doi:10.1007/978-3-642-25847-3_1.

43. Winkler, J. K. et al. Anti-programmed cell death-1 therapy in nonmelanoma skin cancer. British Journal of Dermatology (2017) doi:10.1111/bjd.14664.

44. Ikeda, S. et al. Metastatic basal cell carcinoma with amplification of PD-L1: Exceptional response to anti-PD1 therapy. npj Genomic Med. (2016) doi:10.1038/npjgenmed.2016.37.

45. Falchook, G. S. et al. Responses of metastatic basal cell and cutaneous squamous cell carcinomas to anti-PD1 monoclonal antibody REGN2810. J. Immunother. Cancer (2016) doi:10.1186/s40425-016-0176-3.

46. Terry, S. et al. New insights into the role of EMT in tumor immune escape. Molecular Oncology (2017) doi:10.1002/1878-0261.12093.

47. Proverb, G. B Cell Development, Activation and Effector Functions. Primer to the Immune Response (2014). doi:10.1016/b978-0-12-385245-8.00005-4.

48. Hafemeister, C. & Satija, R. Normalization and variance stabilization of single-cell RNA-seq data using regularized negative binomial regression. Genome Biol. (2019) doi:10.1186/s13059-019-1874-1.

49. Stuart, T. et al. Comprehensive Integration of Single-Cell Data. Cell (2019) doi:10.1016/j.cell.2019.05.031.

50. Kuleshov, M. V. et al. Enrichr: a comprehensive gene set enrichment analysis web server 2016 update. Nucleic Acids Res. (2016) doi:10.1093/nar/gkw377.

51. Murdoch, C., Muthana, M., Coffelt, S. B. & Lewis, C. E. The role of myeloid cells in the promotion of tumour angiogenesis. Nature Reviews Cancer (2008) doi:10.1038/nrc2444.

52. Zhou, P. & Li, T. Construction of the landscape for multi-stable systems: Potential landscape, quasi-potential, A-type integral and beyond. J. Chem. Phys. (2016) doi:10.1063/1.4943096.

53. Wang, J., Zhang, K., Xu, L. & Wang, E. Quantifying the Waddington landscape and biological paths for development and differentiation. Proc. Natl. Acad. Sci. U. S. A. (2011) doi:10.1073/pnas.1017017108.

54. Grafke, T., Schäfer, T. & Vanden-Eijnden, E. Long term effects of small random perturbations on dynamical systems: Theoretical and computational tools. in Fields Institute Communications (2017). doi:10.1007/978-1-4939-6969-2_2.

