## Supplementary material for "Divergent resistance mechanisms to immunotherapy explains response in different skin cancers": SFig 1

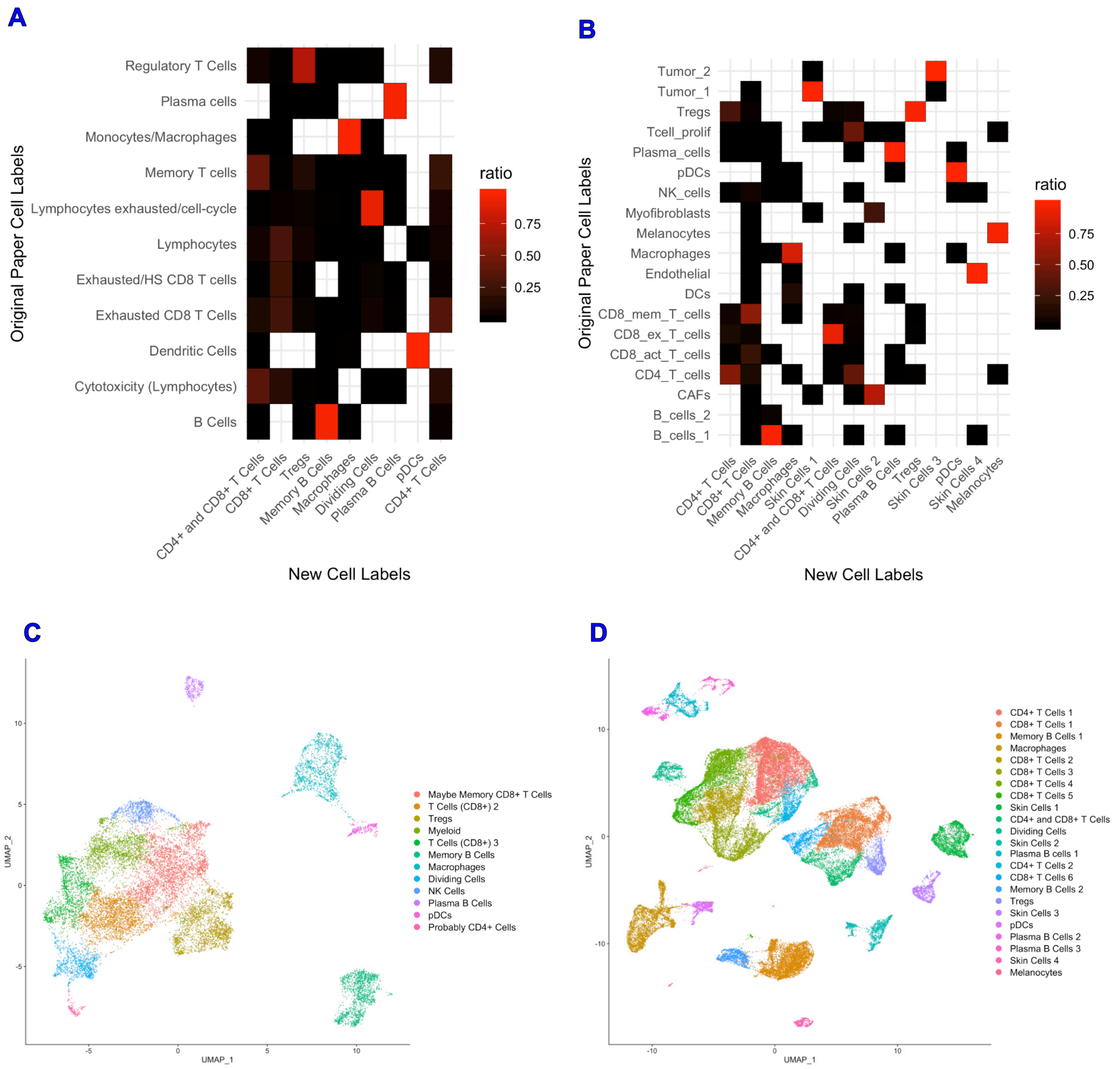


**Supplementary Figure 1: Comparison of cluster identification to original cluster labels.** (A and B): Heatmaps representing the fraction of our clusters in each cluster in the original paper for melanoma (A) and BCC (B). Each column adds to 1. If an original cluster did not contain cells from a new cluster, that space is left blank. (B and D): Dimensionality reduction of the melanoma dataset (C) and the BCC dataset (D), with their original cluster labels. Compare with Fig 1 A and B respectively.
