## Supplementary material for "Divergent resistance mechanisms to immunotherapy explains response in different skin cancers": SFig 2

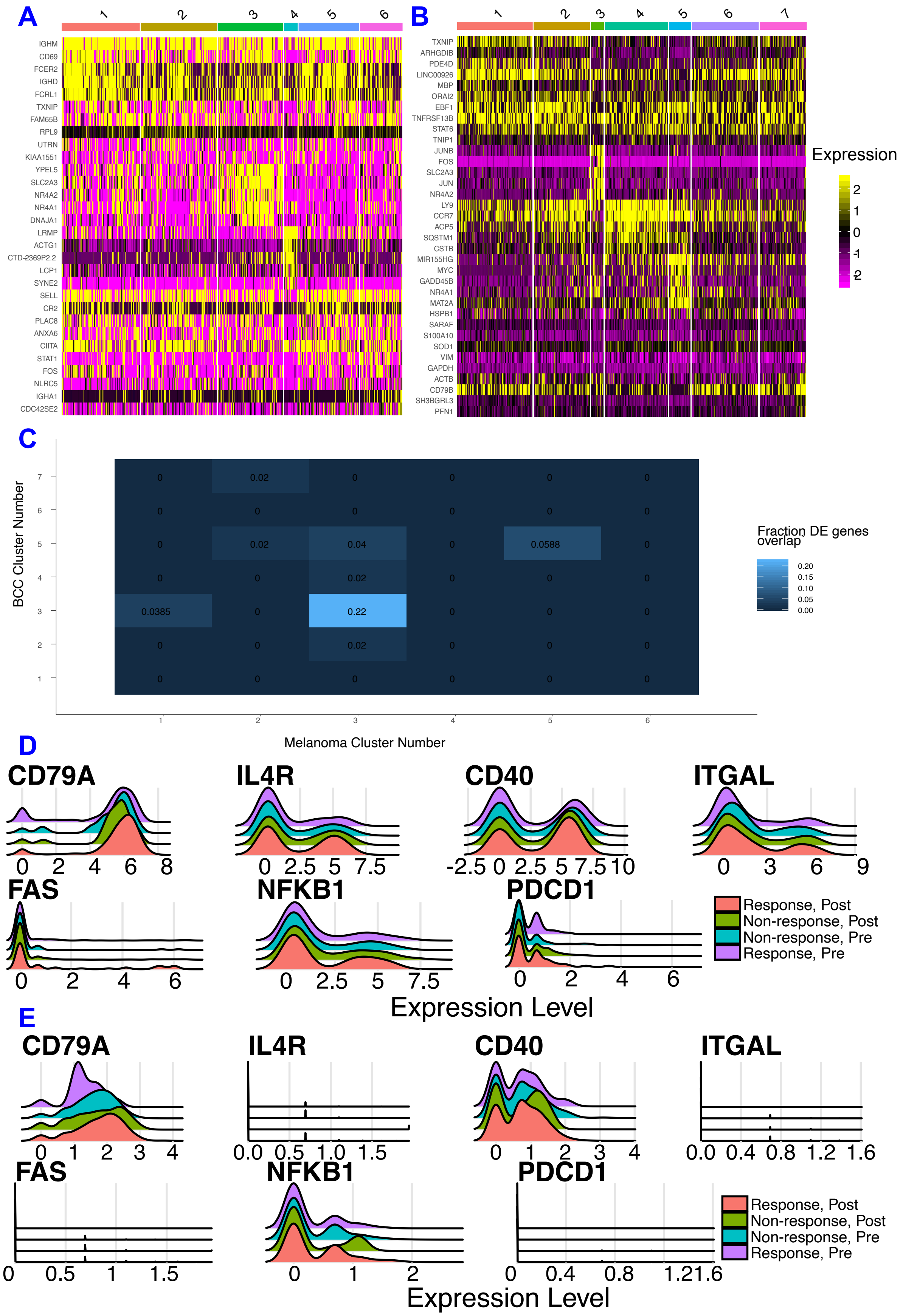
A
1
2
3
4
5
6
IGHM
CD69
FCER2
IGHD
FCRL1
TXNIP
FAM65B
RPL9
UTRN
KIAA1551
YPEL5
SLC2A3
NR4A2
NR4A1
DNAJA1
LRMP
ACTG1
CTD-2369P2.2
LCP1
SYNE2
SELL
CR2
PLAC8
ANXA6
CIITA
STAT1
FOS
NLRC5
IGHA1
CDC42SE2
B
1
2
3
4
5
6
7
TXNIP
ARHGDIB
PDE4D
LINC00926
MBP
ORAI2
EBF1
TNFRSF13B
STAT6
TNIP1
JUNB
FOS
SLC2A3
JUN
NR4A2
LY9
CCR7
ACP5
SQSTM1
CSTB
MIR155HG
MYC
GADD45B
NR4A1
MAT2A
HSPB1
SARAF
S100A10
SOD1
VIM
GAPDH
ACTB
CD79B
SH3BGRL3
PFN1
Expression
2
1
0
-1
-2
C
BCC Cluster Number
Melanoma Cluster Number
Fraction DE genes overlap
0.20
0.15
0.10
0.05
0.00
0.02
0.04
0.0588
0.0385
0.22
D
CD79A
IL4R
CD40
ITGAL
FAS
NFKB1
PDCD1
Response, Post
Non-response, Post
Non-response, Pre
Response, Pre
Expression Level
E
CD79A
IL4R
CD40
ITGAL
FAS
NFKB1
PDCD1
Response, Post
Non-response, Post
Non-response, Pre
Response, Pre
Expression Level

**Supplementary Figure 2: Comparison of memory B cells subsets.** (A and B): Differential expression of memory B cells in melanoma (A) and BCC (B). (C) Fraction of differentially expressed genes in a melanoma and BCC cluster. (D and E): Expression of markers used to calculate the activation and exhaustion score of melanoma (D) and BCC (E).
