## Supplementary material for "Divergent resistance mechanisms to immunotherapy explains response in different skin cancers": SFig 3

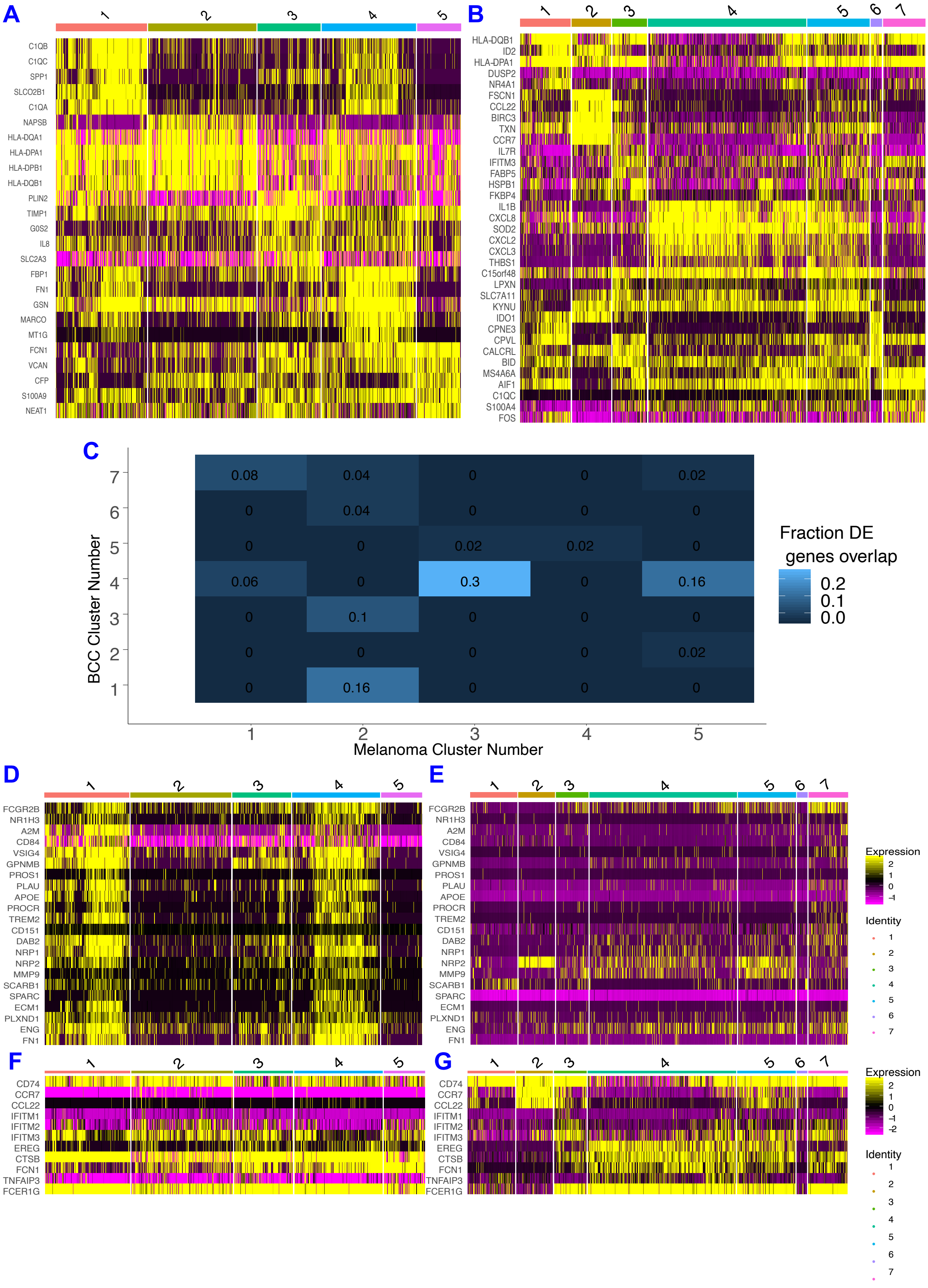
A
C1QB
C1QC
SPP1
SLCO2B1
C1QA
NAPSB
HLA-DQA1
HLA-DPA1
HLA-DPB1
HLA-DQB1
PLIN2
TIMP1
G0S2
IL8
SLC2A3
FBP1
FN1
GSN
MARCO
MT1G
FCN1
VCAN
CFP
S100A9
NEAT1
B
HLA-DQB1
ID2
HLA-DPA1
DUSP2
NR4A1
FSCN1
CCL22
BIRC3
TXN
CCR7
IL7R
IFITM3
FABP5
HSPB1
FKBP4
IL1B
CXCL8
SOD2
CXCL2
CXCL3
THBS1
C15orf48
LPXN
SLC7A11
KYNU
IDO1
CPNE3
CPVL
CALCRL
BID
MS4A6A
AIF1
C1QC
S100A4
FOS
C
BCC Cluster Number
Melanoma Cluster Number
Fraction DE
genes overlap
0.2
0.1
0.0
D
E
FCGR2B
NR1H3
A2M
CD84
VSIG4
GPNMB
PROS1
PLAU
APOE
PROCR
TREM2
CD151
DAB2
NRP1
NRP2
MMP9
SCARB1
SPARC
ECM1
PLXND1
ENG
FN1
Expression
Identity
F
G
CD74
CCR7
CCL22
IFITM1
IFITM2
IFITM3
EREG
CTSB
FCN1
TNFAIP3
FCER1G
Expression
Identity

**Supplementary Figure 3: Comparison of macrophage subsets.** (A and B): Differential expression of the melanoma macrophage clusters (A) and the BCC macrophage clusters (B). (C) Fraction of differentially expressed genes in a melanoma and BCC cluster. (D and E): Expression of anti-inflammatory genes in melanoma macrophages (D) and BCC macrophages (E). (F and G): Expression of pro-inflammatory genes in melanoma macrophages (F) and BCC macrophages (G).
