## Supplementary material for "Divergent resistance mechanisms to immunotherapy explains response in different skin cancers": SFig 4

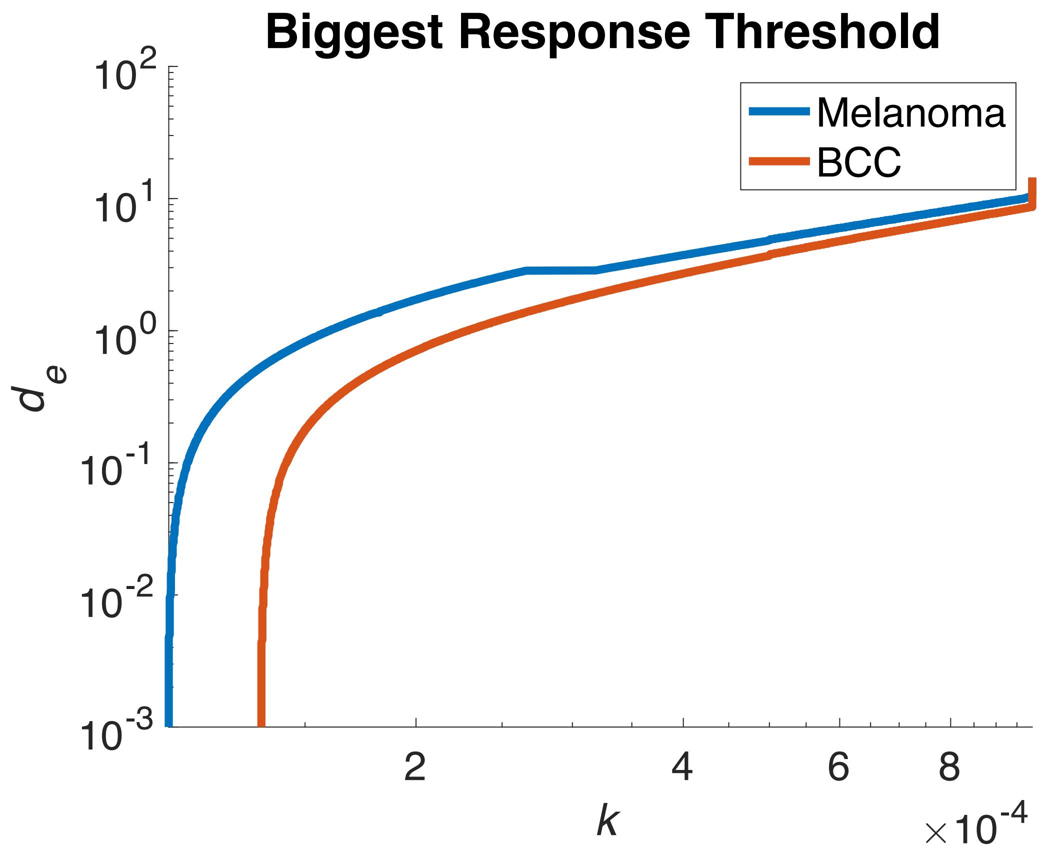


**Supplementary Figure 4: Killing rate at which largest response is observed.** For a fixed $d_e$ value, we determine the $k$ value at which the biggest response occurs. We see that melanoma experiences this maximal response at lower values of $k$ indicating that therapy is more likely to result in a response. By varying $d_e$ along the $y$-axis, we see that this result is robust. To determine the size of the response we sought to maximize, we computed the rate of change of the largest, stable cancer burden as $k$ varied, normalizing for the size of the cancer.
