## Supplementary Tables and Materials for "Divergent resistance mechanisms to immunotherapy explains response in different skin cancers"

| Name | Description | Value | | Units | Source |
|---|---|---|---|---|---|
| | | Melanoma | BCC | | |
| $a$ | Maximum proliferation rate of tumor cells | $0.514$ | | days$^{-1}$ | [1] |
| $b$ | Inverse carrying capacity of tumor | $1.02 \times 10^{-9}$ | | cells$^{-1}$ | [1] |
| $b_e$ | Maximum memory B cell proliferation rate | $1.5$ | | days$^{-1}$ | [2,3] |
| $d$ | Death rate of memory B cells | $2$ | | days$^{-1}$ | [2] |
| $d_e$ | Maximum rate of cancer-mediated deactivation of memory B cells | $1$ | | days$^{-1}$ | Varied as bifurcation parameter |
| $d_m$ | Death rate of macrophages | $3$ | | days$^{-1}$ | Estimated from [4] |
| $g$ | Source rate of macrophages | $30$ | | cells·days$^{-1}$ | Estimated from [4] |
| $k$ | Memory B cell killing rate | $10^{-4}$ | | cells$^{-1}$ · days$^{-1}$ | [1,5–7]; Varied as bifurcation parameter |
| $\kappa_a$ | EC50 for apoptotic-signaling-induced proliferation of macrophages | $4.2 \times 10^{7}$ | $8.4 \times 10^{7}$ | cells·days$^{-1}$ | [8]; Figure 1 |
| $\kappa_d$ | EC50 for cancer-mediated upregulation of memory B cell deactivation | $2.5 \times 10^{6}$ | $2.5 \times 10^{4}$ | cells | Figure 1 |
| $\kappa_e$ | EC50 for cancer-mediated upregulation of memory B cell proliferation | $500$ | | cells | Estimated; [9–11] |
| $\kappa_m$ | EC50 for macrophage-mediated downregulation of memory B cell proliferation | $500$ | $11$ | cells | Estimated |

| | | | | |
|---|---|---|---|---|
| $p$ | Maximum rate of apoptotic-signaling-induced proliferation of macrophages | 4 | days$^{-1}$ | Estimated |
| $s$ | Source rate of memory B cells | $5 \times 10^3$ | cells·days$^{-1}$ | [12,13] |

**Supplementary Table 1: ODE model parameters.**

**Non-dimensionalization of the dynamical model and parameter selection**

We non-dimensionalized our system to simplify the analysis but present our results in terms of these original equations. These are the non-dimensionalized equations:

$$\frac{dx}{d\tau} = x(1-x) - xy$$

$$\frac{dy}{d\tau} = \alpha - \beta y + \gamma \frac{x}{\delta + x}\frac{1}{1+z}y - \epsilon \frac{x}{\zeta + x}y$$

$$\frac{dz}{d\tau} = \eta - \nu z + \theta \frac{xy}{\lambda + \mu xy}z$$

In the table below, $t$ represents the original time variable and τ represents the new time variable.

| | | |
|---|---|---|
| $x = bC$ | $y = \frac{k}{a}B$ | $z = \frac{1}{\kappa_m}M$ |
| $\tau = at$ | $\alpha = \frac{ks}{a^2}$ | $\beta = \frac{d}{a}$ |
| $\gamma = \frac{b_e}{a}$ | $\delta = b\kappa_e$ | $\epsilon = \frac{d_e}{a}$ |
| $\zeta = b\kappa_d$ | $\eta = \frac{g}{a\kappa_m}$ | $\theta = p$ |
| $\lambda = b\kappa_a$ | $\mu = a$ | $\nu = \frac{d_m}{a}$ |

**Supplementary Table 2. Defining relations of non-dimensionalized ODE system.**

**Equilibria and their stability of deterministic model**

In solving for equilibria, we are able to simplify the set of equations into a union of two solution sets:

$$C = 0 \quad \text{or} \quad f(C) = 0$$

where $f$ is a degree 5 polynomial. Of the six roots, we select only the sensical ones, i.e. those where $(C, B, M)$ lies in the first octant of $\mathbb{R}^3$. We determine the stability of these fixed points using the eigenvalues of the Jacobian, looking for those with all eigenvalues having negative real part. In Figure 4B, we show the number of stable equilibria in the $k$-$d_e$ plane. We choose to classify the stable fixed points based on the size of the cancer population. When $C = 0$ is stable, this is complete elimination. When $C$ is nonzero but several orders of magnitude smaller than its carrying capacity, we describe this as a dormant state. When $C$ is at least 1% of the carrying capacity, we call this a high cancer state. When two such states are stable, we call the one with the larger cancer population very high.

**Adding stochastic effect to the model**

We next turned to the reality of biological noise and considered how this could impact our model. Let $X_t$ be the state vector of our system at time $t$ and let $b(X_t)$ be our time-independent ODE function. We take a generic, time-independent noise term, $\sigma(X_t)$ and generated the following SDE model:

$$dX_t = b(X_t)dt + \sigma(X_t)dW_t$$

with $W_t$ being a standard Weiner process. For our noise term, $\sigma$, we assume there are both additive and multiplicative sources of noise for each population. These are given by independent Weiner processes and so $\sigma$ is a $3 \times 6$ matrix. For determining the parameters of these functions, we chose them to be proportional to the parameters of our ODE system (see table below). For B cells and macrophages, these choices were tied to their source rates (additive coefficient) and proliferation rates (multiplicative coefficient). The energy landscapes shown in Figure 5A scaled these noise terms by $1/a$.

| State Variable | Additive Proportionality Constant | Multiplicative Proportionality Constant |
|---|---|---|
| $C$ | $400a$ | $0.4a$ |
| $B$ | $0.04s$ | $0.2b_e$ |
| $M$ | $0.04g$ | $0.2p$ |

**Supplementary Table 3. Noise parameters.** With these parameters, the matrix $\sigma$ is

$$\sigma(C, B, M) = \begin{bmatrix} 400a & 0 & 0 & 0.4a \cdot C & 0 & 0 \\ 0 & 0.04s & 0 & 0 & 0.2b_e \cdot B & 0 \\ 0 & 0 & 0.04g & 0 & 0 & 0.2p \cdot M \end{bmatrix}$$

**Supplementary references:**


1. Pillis, L. G. de, Radunskaya, A. E. & Wiseman, C. L. A Validated Mathematical Model of Cell-Mediated Immune Response to Tumor Growth. *Cancer Res.* **65**, 7950–7958 (2005).
2. Conway, J. M. & Perelson, A. S. A hepatitis C virus infection model with time-varying drug effectiveness: solution and analysis. *PLoS Comput. Biol.* **10**, (2014).
3. Davenport, M. P., Ribeiro, R. M., Chao, D. L. & Perelson, A. S. Predicting the impact of a nonsterilizing vaccine against human immunodeficiency virus. *J. Virol.* **78**, 11340–11351 (2004).
4. Wang, Y. *et al.* Mathematical modeling and stability analysis of macrophage activation in left ventricular remodeling post-myocardial infarction. *BMC Genomics* **13**, S21 (2012).
5. Elemans, M., Florins, A., Willems, L. & Asquith, B. Rates of CTL killing in persistent viral infection in vivo. *PLoS Comput. Biol.* **10**, (2014).
6. Ganusov, V. V *et al.* Fitness costs and diversity of the cytotoxic T lymphocyte (CTL) response determine the rate of CTL escape during acute and chronic phases of HIV infection. *J. Virol.* **85**, 10518–10528 (2011).
7. Wick, W. D., Yang, O. O., Corey, L. & Self, S. G. How many human immunodeficiency virus type 1-infected target cells can a cytotoxic T-lymphocyte kill? *J. Virol.* **79**, 13579–13586 (2005).
8. Murdoch, C., Muthana, M., Coffelt, S. B. & Lewis, C. E. The role of myeloid cells in the promotion of tumour angiogenesis. *Nature Reviews Cancer* (2008) doi:10.1038/nrc2444.
9. Cabrita, R. *et al.* Tertiary lymphoid structures improve immunotherapy and survival in melanoma. *Nature* 1–5 (2020).
10. Petitprez, F. *et al.* B cells are associated with survival and immunotherapy response in sarcoma. *Nature* **577**, 556–560 (2020).
11. Helmink, B. A. *et al.* B cells and tertiary lymphoid structures promote immunotherapy response. *Nature* **577**, 549–555 (2020).
12. Althaus, C. L. & De Boer, R. J. Dynamics of immune escape during HIV/SIV infection. *PLoS Comput. Biol.* **4**, (2008).
13. De Boer, R. J. Understanding the failure of CD8+ T-cell vaccination against simian/human immunodeficiency virus. *J. Virol.* **81**, 2838–2848 (2007).
